# PIC50: An open source tool for interconversion of PIC_50_ values and IC_50_ for efficient data representation and analysis

**DOI:** 10.1101/2022.10.15.512366

**Authors:** Aman Thakur, Ajay Kumar, Vivek Sharma, Vineet Mehta

## Abstract

Half-maximal inhibitory concentration (IC_50_) is used to determine the potency of a drug against a variety of enzymes/ biological targets associated with the pathogenesis of multiple disorders. The IC_50_ values can be depicted in multiple ways, which makes it difficult to analyze the results presented in different concentrations. Representing data in the form of PIC_50_ values depicting the IC_50_ values as the negative logarithm of IC_50_ in molar concentration is considered to be a better approach as it not only makes data easily understandable but also eliminates the possibility of errors in data representation and reproducibility. Considering the importance of data representation for a better understanding of data and comparing efficacy and potency of the drugs, besides, the significance of PIC_50_ value in the field of CADD, we found that at present there is no single open-source software available to convert the IC_50_ values to PIC_50_ values and vice versa from millimolar to picomolar range. Therefore, in the present study, we develop a tool that could help researchers to interconvert IC_50_ values and PIC_50_ values in a reliable way to eliminate the possibility of errors. We validated our tool through three case studies where the data generated by our tool was found to be 100% accurate. Moreover, we present a case where data was published in literature with errors in calculated PIC_50_ values and demonstrated the importance and reliability of our tool.

## Introduction

Half-maximal inhibitory concentration (IC_50_) is one of the extensively acknowledged parameters to assess the potency of a drug against a variety of enzymes/ biological targets that are associated with the pathogenesis of disorders like Alzheimer’s disease, depression, diabetes mellitus, cancer, etc. (Aykul and Martinez-Hackert, et al., 2016). IC_50_ is a measure of the potency of a drug and is defined as the minimum concentration obligatory for inhibiting the biological activity of a target by 50% (Aykul and Martinez-Hackert, et al., 2016). Literature reports suggest that the IC_50_ values of a particular drug against a particular enzymes/ biological targets can be depicted in multiple ways, which makes it difficult to analyze the results and reach a decisive conclusion regarding the exact efficacy and potency of a drug (Changyong et al., 2014). One of the better ways to present data of a scientific finding is in a logarithmic form as it spreads out the data in such a way that the shape of the curve and the quality of the fit are readily visible when the concentrations cover a wide range. There is a group of scientists that argue that linear functions can be used for percent inhibition but should not be used to calculate IC_50_ as the IC_50_ curve is not a linear function, rather it is a saturating function. Representing data in the form of PIC_50_ values is considered to be a better approach as it not only makes data easily understandable but also eliminates the possibility of errors in data representation and reproducibility (Changyong et al., 2014). PIC_50_ is the approach for depicting the IC_50_ values as the negative logarithm of IC_50_ in molar concentration, therefore it makes data more convenient for the readers to understand and compare the potency of different drugs at the same molar levels (Abdulrahman et al., 2021; Thakur et al., 2022). PIC_50_ values are now being extensively used in an array of computer-aided drug designing (CADD) approaches such as Quantitative Structural Activity Relationship (QSAR) (Hadaji et al., 2017; Ramalakshmi et al., 2021), comparative molecular field analysis (coMFA), comparative molecular similarity indices analysis (coMSIA) (Liang et al., 2013), Pharmacophore modeling (Vyas et al., 2013), etc. and the outcome of these tools solely depends on the reliability of input data of PIC_50_ values. As the IC_50_ value is distributed on a wider range, hence to alter these values to a normal distribution, the PIC_50_ value is used so that a vast numerical distribution can be represented in certain intervals (da Silva Costa et al., 2018; Ferreira et al., 2019). Moreover, it also endows with a logarithmic approach to the study as higher values of PIC_50_ signify higher potency (Hendrickx et al., 2018). Although the approach to convert IC_50_ values to PIC_50_ values seems to be simple, through rigorous literature review we found that it is a tricky process and there are significant errors in the calculated PIC_50_ values in published research papers.

Considering the importance of data representation for a better understanding of data and comparing efficacy and potency of the drugs, besides, the significance of PIC_50_ value in the field of CADD, we found that at present there is no single open-source software available to convert the IC_50_ values to PIC_50_ values and vice versa from milimolar to picomolar range. Therefore, in the present study, we aimed to develop a tool that can help researchers to interconvert IC_50_ values and PIC_50_ values in a reliable way to eliminate the possibility of errors. This software will not only reduce the time for calculating the PIC_50_ or IC_50_ values but significantly eliminates the chance of human error in calculation.

## Material and Methods

### Developed PIC50 Tool

The current PIC_50_ tool can be downloaded free from https://www.researchgate.net/publication/363769944_PIC50_to_IC50_and_IC50_to_PIC50_calculator#fullTextFileContent. This tool allows user to calculate PIC_50_ values form the IC_50_ values depicted in millimolar, micromolar, nanomolar, or picomolar concentration range.

### Software and its functionalities

The current software “PIC50” is developed on python programming language and further converted to .exe file format for ease of portability. It works at ease on and above Windows 7 platforms. Currently, it consists of two modules: (i) Calculating IC_50_ from PIC_50_ value and (ii) Calculating PIC_50_ from IC_50_ value.

After opening the .exe file, the user will get two options for calculating IC_50_ or PIC_50_, enter “1” for calculating the IC_50_ value from PIC_50_ and “2” for calculating the PIC_50_ value from IC_50_ value. Afterward, the user will get further options to calculate their values either in millimolar, micromolar, nanomolar, or picomolar range as per the requirement. Users will be prompted to enter “3” for millimolar, “6” for micromolar, “9” for nanomolar, and “12” for picomolar values. After entering the IC_50_/PIC_50_ values user will simply get their respective PIC_50_/IC_50_ values (Image 1). There is no limit for calculating the number of values as a user can calculate values ‘n’ times from the software. The current software uses the following equations for calculating the IC_50_ and PIC_50_ values respectively:

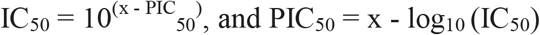

**Image 1:**
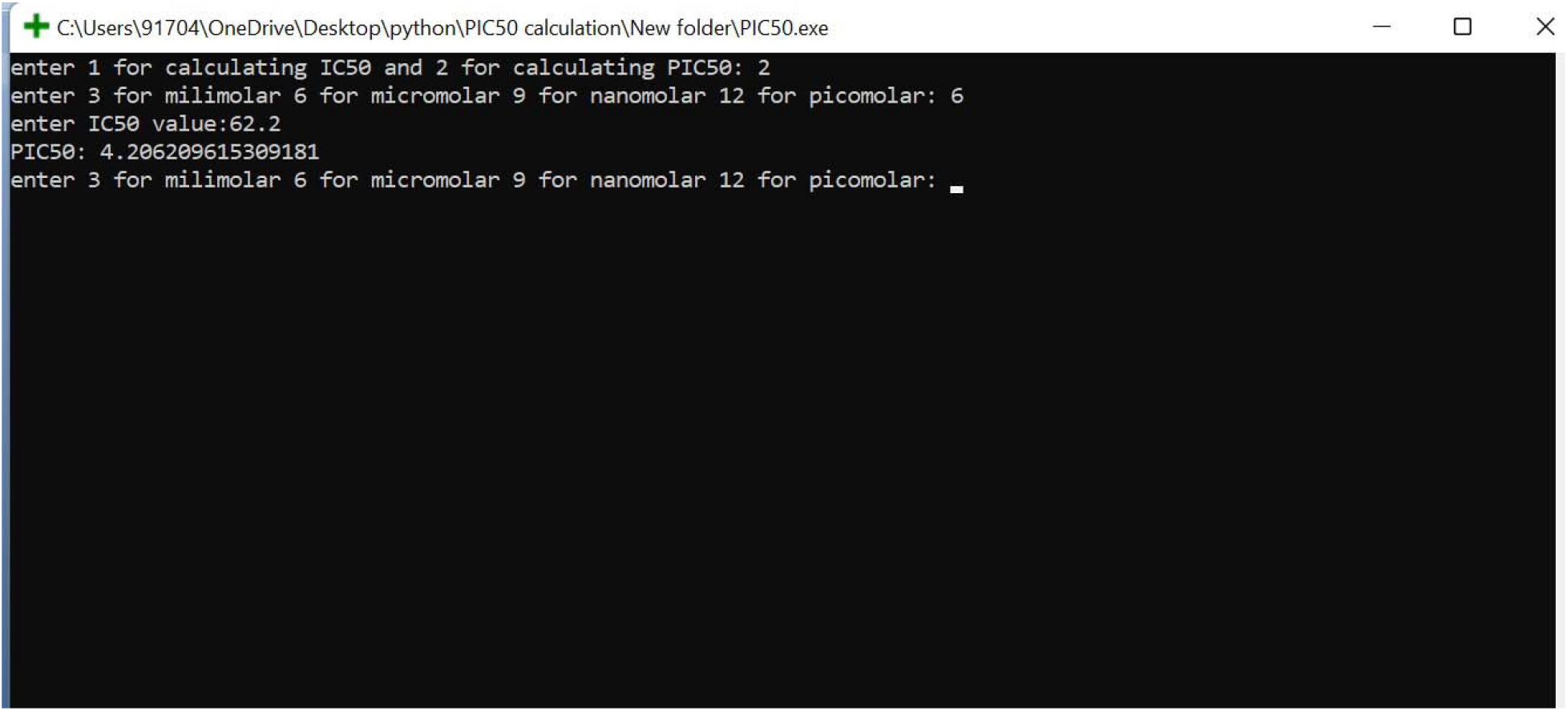
Picture showing the calculation of PIC_50_ in the micromolar range.

Where, x = 3 for millimolar, 6 for micromolar, 9 for nanomolar, and 12 for picomolar concentrations

## Results and Discussion

To confirm the functionality of the current software we performed three case studies. In the first case study, a series of 30 1H□Pyrazole-1-carbothioamide derivatives were employed to develop a QSAR model, and based on this model PIC_50_ value (in micromolar) of novel 11 derivatives was calculated. In our study, we predicted accurately the PIC_50_ of 30 derivatives used and also converted the PIC_50_ to IC_50_ of 11 compounds, whose activity was predicted for comparative study with the 30 derivatives (Hajalsiddig et al., 2020). Results are depicted in Table 1 (a, b). In this study, author calculated experimental PIC_50_ value of compound “C16” as 5.24 whereas correct experimental PIC_50_ value of the same is calculated from our tool is 5.27. This error in calculating PIC_50_ value of this compound also resulted in wrong interpretation of residuals (Experimental PIC_50_; Predicted PIC_50_) from the two QSAR equations generated in the study. In the equation 1, author has calculated residual value of compound “C16” as 0.12, however, it should be 0.15. Likewise, in equation 2, the value of the same was reported to be 0.9 but it should be 0.12. This error will be automatically be carried forward in the whole QSAR model generation and will affect the overall accuracy of the QSAR model significantly.

**Table 1a:** Reported PIC_50_ values of the 30 compounds (Hajalsiddig et al., 2020) and PIC_50_ values predicted by our tool.

| S. No. | Name of Compound | PIC<sub>50</sub> calculated by the authors | PIC<sub>50</sub> calculated by our tool |
| --- | --- | --- | --- |
| 1 | C1 | 6.08 | 6.081 |
| 2 | C2 | 5.87 | 5.866 |
| 3 | C3 | 5.66 | 5.665 |
| 4 | C4 | 6.47 | 6.468 |
| 5 | C5 | 7.15 | 7.155 |
| 6 | C6 | 6.89 | 6.886 |
| 7 | C7 | 5.51 | 5.514 |
| 8 | C8 | 5.28 | 5.285 |
| 9 | C9 | 5.41 | 5.412 |
| 10 | C10 | 5.37 | 5.375 |
| 11 | C11 | 5.20 | 5.203 |
| 12 | C12 | 5.14 | 5.135 |
| 13 | C13 | 5.16 | 5.164 |
| 14 | C14 | 5.13 | 5.132 |
| 15 | C15 | 5.24 | 5.241 |
| **16** | **C16** | **5.24** | **5.278** |
| 17 | C17 | 5.11 | 5.109 |
| 18 | C18 | 5.18 | 5.183 |
| 19 | C19 | 5.13 | 5.137 |
| 20 | C20 | 5.19 | 5.191 |
| 21 | C21 | 5.04 | 5.050 |
| 22 | C22 | 5.08 | 5.089 |
| 23 | C23 | 4.99 | 4.997 |
| 24 | C24 | 5.00 | 5.007 |
| 25 | C25 | 5.09 | 5.092 |
| 26 | C26 | 4.95 | 4.948 |
| 27 | C27 | 4.97 | 4.973 |
| 28 | C28 | 4.87 | 4.874 |
| 29 | C29 | 4.91 | 4.911 |
| 30 | C30 | 4.95 | 4.960 |

**Table 1b:**
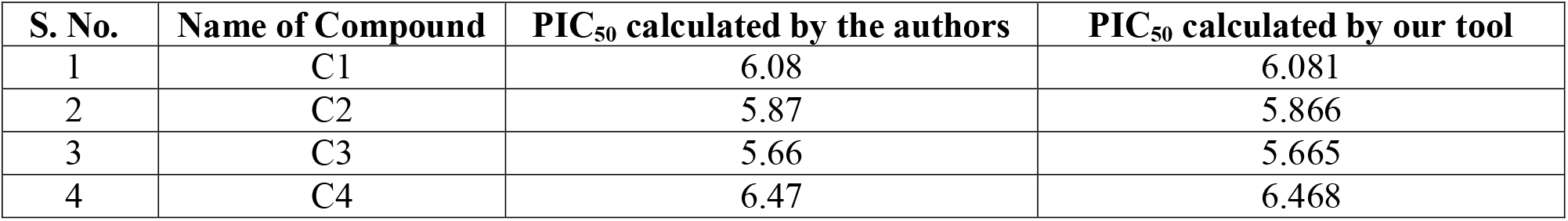
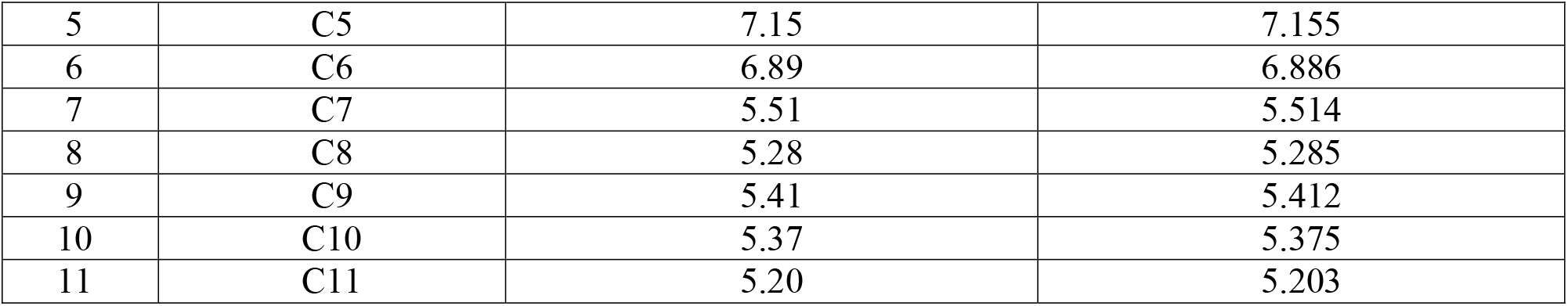
Reported PIC_50_ values of the 11 compounds (Hajalsiddig et al., 2020) and PIC_50_ values predicted by our tool.

| S. No. | Name of Compound | PIC<sub>50</sub> calculated by the authors | PIC<sub>50</sub> calculated by our tool |
| --- | --- | --- | --- |
| 1 | C1 | 6.08 | 6.081 |
| 2 | C2 | 5.87 | 5.866 |
| 3 | C3 | 5.66 | 5.665 |
| 4 | C4 | 6.47 | 6.468 |
| 5 | C5 | 7.15 | 7.155 |
| 6 | C6 | 6.89 | 6.886 |
| 7 | C7 | 5.51 | 5.514 |
| 8 | C8 | 5.28 | 5.285 |
| 9 | C9 | 5.41 | 5.412 |
| 10 | C10 | 5.37 | 5.375 |
| 11 | C11 | 5.20 | 5.203 |

In the second study, 28 berberine analogs were used to develop a QSAR model against coxsackie B1 virus and PIC_50_ values of all the analogs were measured in micromolar. We also used our software and predicted the PIC_50_ values accurately and in a very short time period (Obadawo et al., 2022). Results are depicted in Table 2. In this study, author has calculated the PIC_50_ value of only 24 compounds out of 28 compounds mentioned in the literature. The PIC_50_ values of Compound No. “4”, “19”, “26” and “27” was not calculated by the author. Our tool successfully calculated the PIC_50_ of these excluded compounds accurately. This suggest that the tool developed in the present study is efficient to calculate PIC_50_ values over wide range of concentrations reliably and accurately.

**Table 2:** Reported PIC_50_ values of the 28 compounds (Obadawo et al., 2022) and PIC_50_ values predicted by our tool.

| S. No. | Name of Compound | $PIC_{50}$ calculated by author | $PIC_{50}$ calculated by our tool |
| --- | --- | --- | --- |
| 1 | 1 | 5.644 | 5.644 |
| 2 | 2 | 5.4225 | 5.422 |
| 3 | 3 | 5.2628 | 5.262 |
| **4** | **4** | **Not Calculated by author** | **5.634** |
| 5 | 5 | 4.9066 | 4.907 |
| 6 | 6 | 4.8665 | 4.866 |
| 7 | 7 | 5.1918 | 5.192 |
| 8 | 8 | 5.2984 | 5.298 |
| 9 | 9 | 5.5058 | 5.506 |
| 10 | 10 | 5.5918 | 5.592 |
| 11 | 11 | 4.8665 | 4.866 |
| 12 | 12 | 5.2097 | 5.210 |
| 13 | 13 | 4.983 | 4.983 |
| 14 | 14 | 5.2182 | 5.218 |
| 15 | 15 | 5.3019 | 5.302 |
| **16** | **16** | **5.0353** | **5.035** |
| 17 | 17 | 4.6799 | 4.680 |
| 18 | 18 | 4.9626 | 4.963 |
| **19** | **19** | **Not calculated by the author** | **4.487** |
| 20 | 20 | 4.3925 | 4.392 |
| 21 | 21 | 5.0353 | 5.035 |
| 22 | 22 | 4.9066 | 4.907 |
| 23 | 23 | 4.5406 | 4.541 |
| 24 | 24 | 4.8268 | 4.827 |
| 25 | 25 | 5.1681 | 5.168 |
| **26** | **26** | **Not calculated by the author** | **4.866** |
| **27** | **27** | **Not calculated by the author** | **4.801** |
| 28 | 28 | 5.1662 | 5.166 |

In the third case, 18 compounds were employed for developing a QSAR model for predicting anti-malarial activity. The PIC_50_ values (in the nanomolar range) were used for performing the QSAR study. The same was predicted by our software accurately (Sahu et al., 2014). Results are depicted in Table 3. In this study two QSAR models were generated, one against Chloroquine sensitive *Plasmodioum falciparum* strain (HB3) and second for Chloroquine resistant *Plasmodioum falciparum* strain (Dd2). In the PIC_50_ value (in nanomolar range), author has calculated the PIC_50_ value of compound “16b” as 7.119, however, the accurate value of the same should be 7.286 that was predicted accurately by the tool developed in present study. Moreover, for compound “17a” the PIC_50_ value calculated by author was 7.286 whereas the same should be 7.119, as calculated by our tool. This error will be automatically be carried forward in the whole QSAR model generation and will affect the overall accuracy of the QSAR model significantly.

**Table 3:** Reported PIC_50_ values of the 18 compounds (Sahu et al., 2014) and PIC_50_ values predicted by our tool.

| S. No. | Name Of Compound | PIC<sub>50</sub> calculated by author | PIC<sub>50</sub> calculated by our tool |
| --- | --- | --- | --- |
| 1 | 4a | 6.889 | 6.889 |
| 2 | 4b | 7.249 | 7.249 |
| 3 | 4c | 6.77 | 6.770 |
| **4** | 4d | 6.987 | 6.987 |
| 5 | 4e | 6.57 | 6.570 |
| 6 | 5a | 7.506 | 7.506 |
| 7 | 5b | 7.551 | 7.551 |
| 8 | 5c | 7.073 | 7.073 |
| 9 | 5d | 7.363 | 7.363 |
| 10 | 5e | 6.562 | 6.562 |
| 11 | 6a | 6.893 | 6.893 |
| 12 | 6b | 7.001 | 7.001 |
| 13 | 7a | 6.055 | 6.055 |
| 14 | 7b | 5.593 | 5.593 |
| 15 | 16a | 7.097 | 7.097 |
| **16** | **16b** | **7.119** | **7.286** |
| 17 | **17a** | **7.286** | **7.119** |
| 18 | 17b | 7.121 | 7.121 |

In another case study, a total of twenty-one 3-Iodochrome derivatives were synthesized and the QSAR model was generated for potential fungicides (Kaushik et al., 2021). This study is a perfect example of incorrect data representation arising due to a lack of a proper tool to calculate PIC_50_ values from the IC_50_ values. In this study, ED_50_ values of the compounds were evaluated against *Sclerotium rolfsii* in the mgL^-1^ (Kaushik et al., 2021). For developing the QSAR model, the author utilized the pED_50_ values, however, for these calculations authors did not consider the factor of the unit of measurement i.e. mg (10^−3^). We calculated the PIC_50_ values from our software and compared them to the reported values. Table 4 depicts the pED_50_ value calculated by the author (Kaushik et al., 2021) and the pED_50_ value calculated by our software from the given ED_50_.

**Table 4:** Difference in the reported pED_50_ and pED_50_ values calculated by our software.

| S. No. | Compound Name | $ED_{50}$ ($mgL^{-1}$) | $pED_{50}$<br>(Calculated by the author) | $pED_{50}$<br>(Calculated by our software) |
| --- | --- | --- | --- | --- |
| 1 | 4a | 50.38 | -1.70 | 1.30 |
| 2 | 4b | 19.12 | -1.28 | 1.72 |
| 3 | 4c | 30.2 | -1.48 | 1.52 |
| 4 | 4d | 39.28 | -1.59 | 1.41 |
| 5 | 4e | 47.59 | -1.68 | 1.32 |
| 6 | 4f | 61.23 | -1.79 | 1.21 |
| 7 | 4g | 64.02 | -1.81 | 1.19 |
| 8 | 4h | 75.58 | -1.88 | 1.12 |
| 9 | 4i | 93.17 | -1.97 | 1.03 |
| 10 | 4j | 124.7 | -2.10 | 0.90 |
| 11 | 4k | 165.7 | -2.22 | 0.78 |
| 12 | 4l | 248.6 | -2.40 | 0.60 |
| 13 | 4m | 325 | -2.51 | 0.49 |
| 14 | 4n | 410 | -2.61 | 0.39 |
| 15 | 4o | 515.4 | -2.71 | 0.29 |
| 16 | 4p | 72.86 | -1.86 | 1.14 |
| 17 | 4q | 63.75 | -1.80 | 1.20 |
| 18 | 4r | 8.43 | -0.93 | 2.07 |
| 19 | 4s | 147.8 | -2.17 | 0.83 |
| 20 | 4t | 33.98 | -1.53 | 1.47 |
| 21 | 4u | 110.9 | -2.04 | 0.96 |

Considering the example of compound ‘4a’ from the table, ED_50_ was reported to be 50.38 mgL^-1^. The authors computed the negative logarithm of ED_50_ values directly of the compound ‘4a’ without taking the 10^−3^ (for milligram) into the calculation. Since pED_50_ values are the negative logarithm of EC_50_ values in molar concentration, the concentration factor has to be taken into consideration. From our tool, pED_50_ of the same molecules was calculated by taking the unit of milligram also into consideration and the values are depicted in Table 4. Interestingly, our results demonstrated significant variations from the reported values. Our software utilized the following calculation for calculating the pED_50_ values:

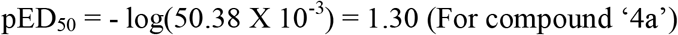

It is evident from Table 4 that our tool calculated the pED_50_ values accurately and also solved the purpose of using the pED_50_ value, which is the logarithmic approach (higher values for more potent compounds).

From our study, we efficiently calculated PIC_50_ values from IC_50_ values for a total of 108 compounds that were randomly selected from literature. We also identified the error in data presentations in these studies, which was carried forward in the whole QSAR model generation and had adversely affected the overall accuracy of the QSAR model significantly. Moreover, in the last case study, entire calculations were found to have error in the determination of PIC_50_ values, which were predicted accurately by our tool.

## Conclusion

QSAR is a well-established technique that has a wide application in the field of drug designing, pharmaceuticals, ecotoxicity of industrial chemicals, materials science, etc. (Puzyn et al., 2010; Roy et al., 2017). The accurate depiction of the PIC_50_ values is a crucial step for developing robust and reliable QSAR models. Currently, there is no tool available that can accurately and reliably calculate the PIC50 values and there are significant chances of errors in manual methods. In our study, we have presented a user-friendly, simple, and convenient PIC_50_ value calculator that can accurately calculate PIC_50_ values to eliminate any possibility of errors and present error-free data in an easily understandable and reliable manner. This software is available in open source (https://www.researchgate.net/publication/363769944_PIC50_to_IC50_and_IC50_to_PIC50_calculator/stats) and can be easily downloaded and used. Our software not only calculates the PIC_50_ value from the IC_50_ value but also performs the IC_50_ calculations from the PIC_50_ value. The developed PIC_50_ tool is capable of calculating values in all ranges of concentrations including millimolar, micromolar, nanomolar, and picomolar, which makes this tool more valuable. Further, to evaluate the functionality of our tool we examined three case studies and found all the PIC_50_ and IC_50_ values were calculated with 100% accuracy and in very less time. Moreover, our results (Table 1 to Table 4) demonstrate how there are vast possibilities of errors in data representation and how that can be eliminated by using the developed PIC_50_ tool. For any research, the accuracy of the data and time saving are the two most important criteria. Our software provides both for all the researchers who used these parameters for their studies on an open-source platform.

## Acknowledgement

The authors acknowledge Govt. College of Pharmacy, Rohru and Institute of Pharmaceutical Science, Kurukshetra University, Haryana for providing the facilities to conduct this study.

## Author’s Contribution

AT contributed to designing the study, performed QSAR, and prepared the first draft of the manuscript. AK and VS contributed to the study designing, technical inputs, and manuscript editing. VM contributed to designing the study, Results analysis, manuscript preparation, and editing analysis of the results.

